# Reproducible and open-source statistical shape models of the pose-normalized adult lower body: development and comparative evaluation

**DOI:** 10.64898/2026.11.21.768300

**Authors:** Muhammad Abdullah, Abdul Aziz Hulleck, Kinda Khalaf, Marwan El-Rich

**Author notes:** Corresponding Author: Marwan El-Rich.

## Abstract

Accurate skeletal geometry is important for subject-specific musculoskeletal modeling, but the choice of statistical representation may affect anatomical reconstruction. This study developed a reproducible workflow and compared four statistical shape model families of the pose-normalized adult lower body: pooled global, articulated shape-pose, sex-conditioned shared-basis, and independently fitted sex-specific models. Models were developed from computed-tomography-derived geometries of 20 adults and evaluated using leave-one-subject-out cross-validation. Model coefficients were estimated from each held-out subject’s complete registered surfaces, assessing dense-surface representational generalization. Outcomes included model compactness, whole-body and bone-wise surface root-mean-square errors, and reconstruction errors for hip, knee, and ankle joint centers. A pelvis-referenced bounding-box anisotropic-scaling baseline was included as a geometric comparator. The pooled global and sex-conditioned models shared a principal-component basis and therefore identical compactness spectra, with the first component explaining 90.16% of the variance. These models provided the most consistent reconstruction performance. The articulated model had ankle and knee joint-center errors 9.1 and 5.4 mm higher, respectively, than the pooled global model. Independently fitted sex-specific models did not improve whole-body reconstruction over the pooled global or sex-conditioned models in this small cohort. Under the present evaluation, the pooled global and sex-conditioned models reduced whole-body surface errors by approximately 4.1 and 4.0 mm, respectively, relative to scaling. The pooled global model also reduced ankle and knee joint-center errors by 12.5 and 11.0 mm. These results characterize the implemented representations within the available dataset. The scaling comparison was not input matched, and sparse-input and external validation remain necessary. The accompanying LowerBodySSM workflow and interactive Companion provide a reproducible resource for further anatomical modeling and evaluation.

## 1. Introduction

Musculoskeletal (MSK) models have become essential tools in biomechanics, sports science, rehabilitation, and engineering because they enable the estimation of internal quantities that are difficult or impossible to measure directly, including joint loading, muscle forces, and the mechanical consequences of hypothetical interventions [1, 2]. However, the utility of these models depends strongly on how accurately a generic model is personalized to an individual’s anatomy. Model personalization or scaling is therefore a central determinant of the anatomical and biomechanical validity of subject-specific simulations [3, 4].

The most widely used personalization approaches scale generic MSK models using subject anthropometrics, segment lengths, body mass, or marker positions from static or dynamic motion-capture trials [5–9]. These linear scaling methods are robust, computationally efficient, and easy to implement, which explains their continued use in large biomechanical studies [10]. Nevertheless, linear scaling assumes that anatomical differences can be represented by uniform or axis-wise changes in size. This assumption cannot fully capture nonlinear anatomical variation such as bone curvature, joint-surface morphology, torsion, relative bone placement, and local attachment-site geometry, all of which can affect joint-center location, muscle paths, moment arms, and predicted loading [11].

Image-based personalization using computed tomography (CT) or magnetic resonance imaging (MRI) can overcome many of these limitations by directly deriving subject-specific bone geometry from medical images and morphing generic models via surface registration or radial basis function approaches [12–14]. Related computational frameworks have similarly emphasized the importance of image-derived geometry and mesh processing for constructing patient-specific models suitable for numerical simulation [15, 16]. Although these methods can provide highly individualized geometry, they require image acquisition, segmentation, mesh processing, and specialized expertise. Their cost, time demand, and operational complexity limit their use in settings where rapid, accessible, or low-burden model personalization is required [7, 17].

Statistical shape models (SSMs) provide an intermediate strategy between simple linear scaling and fully image-based personalization. Their construction depends on establishing meaningful correspondence across the training geometries, since correspondence provides a consistent representation of homologous shape variation across subjects [18]. An SSM learns a population mean shape and principal modes of anatomical variation from a training set of aligned bone meshes. Once trained, the model can reconstruct subject-specific anatomy from sparse landmarks, motion-capture markers, anthropometric variables, or other reduced input descriptions [19–21]. Because SSMs capture coordinated patterns of anatomical variation, they can estimate features that are not recoverable by length-based scaling alone, including joint centers, local bone morphology, and muscle attachment or path-point regions [22, 23].

Early lower-limb SSM work established the feasibility of using population-derived shape variation to generate subject-specific skeletal geometry for MSK model personalization. Zhang et al. [19] introduced the Musculoskeletal Atlas Project (MAP) Client as an open workflow environment for MSK modeling, providing a practical platform for integrating anatomical modeling, registration, and model-scaling tools. Building on this workflow, Zhang et al. [24] developed an articulated lower-limb SSM in which component bones were represented within a linked anatomical framework and reconstructed from sparse anatomical landmarks. Their model included the pelvis, unilateral femur, patella, tibia, and fibula, together with embedded anatomical landmarks and muscle attachment regions for MSK model scaling and personalization. Subsequent MAP Client applications showed that SSM-based scaling could improve hip joint-center (HJC) estimation and muscle moment-arm predictions in people with hip osteoarthritis [23], reduce reliance on segment markers without substantially altering inverse-kinematics outputs [25], and provide greater robustness to marker-placement error than conventional linear scaling [26]. However, these models generally focused on the pelvis and major long bones and did not provide complete bilateral lower-limb coverage, including the distal hindfoot and foot-related domains.

Pediatric lower-limb SSMs have also been developed separately to address population-specific skeletal morphology. This is important because adult-derived SSMs may not generalize well to children, particularly when skeletal size, proportions, and maturation-related shape variation differ substantially from those of adults [27]. Carman et al. [28] introduced an articulated pediatric lower-limb shape model that predicted bone geometry from sparse landmarks, while Carman et al. [29] evaluated regression, linear scaling, and SSM approaches for predicting the pediatric HJC. Shi et al. [30] similarly developed predictive pediatric lower-limb bone models. These studies demonstrate the value of population-specific SSMs, particularly when adult generic scaling is inappropriate. However, their anatomical scope has generally remained focused on the pelvis and major long bones, and their objectives differ from developing a broadly inclusive adult lower-body SSM extending from the pelvis to the metatarsal-containing foot segments.

More recent work has extended SSM-based reconstruction toward broader lower-limb personalization. Zhai et al. [17] proposed a marker-driven SSM framework for reconstructing lower-limb skeletal geometry, including the pelvis and bilateral femur, patella, tibia, and fibula. Audenaert et al. [31] addressed another important modeling challenge by separating positional noise from neutral alignment in multicomponent SSMs, emphasizing that multi-bone models must distinguish true anatomical shape variation from residual pose and alignment effects. These studies represent important progress toward more comprehensive and interpretable lower-limb models. Nevertheless, distal foot structures, such as the talus, calcaneus, and associated foot domains, remain underrepresented, and the influence of alternative SSM formulations on reconstruction accuracy has not been systematically evaluated.

Further evaluation is needed to understand how alternative statistical representations affect reconstruction of multicomponent lower-body anatomy. Pooled global, articulated shape–pose, sex-conditioned shared-basis, and independently fitted sex-specific models differ in how they represent anatomical shape, relative bone placement, and sex-related variation. Comparing these formulations within a common dataset and evaluation framework can clarify their reconstruction performance and the trade-offs associated with separate subgroup models. Broader anatomical coverage, including the talus and metatarsal-containing foot segments, and accessible computational workflows can also support reproducibility, reuse, and further methodological comparisons.

This study developed and evaluated LowerBodySSM, a reproducible framework for statistical shape modeling of the pose-normalized adult lower body using geometries from 20 adults. The models extended from the pelvis and sacrum through the bilateral lower limbs to the metatarsal-containing foot segments, excluding the toes. Four model families were compared in terms of compactness, whole-body and bone-wise surface reconstruction errors, and reconstruction of hip, knee, and ankle joint centers and hip joint-center width. Leave-one-subject-out (LOSO) evaluation used each held-out subject’s complete registered surfaces to estimate model coefficients, thereby assessing dense-surface representational generalization. A study-specific pelvis-referenced bounding-box anisotropic-scaling baseline provided a geometric reference; this comparison was not a matched-input evaluation of clinical personalization. It was hypothesized that the pooled global and sex-conditioned shared-basis models would provide the most consistent reconstruction performance and that the statistical shape models would produce lower errors than the selected scaling baseline. The accompanying workflow, model datasets, validation outputs, and interactive Companion provide a resource for reuse and further evaluation.

## 2. Materials and Methods

The study workflow is summarized in Fig. 1. It comprised anatomical alignment and pose normalization, dense mesh correspondence, statistical shape model construction, and LOSO dense-surface reconstruction and comparison. Each step is described in the following subsections.

**Fig. 1.**
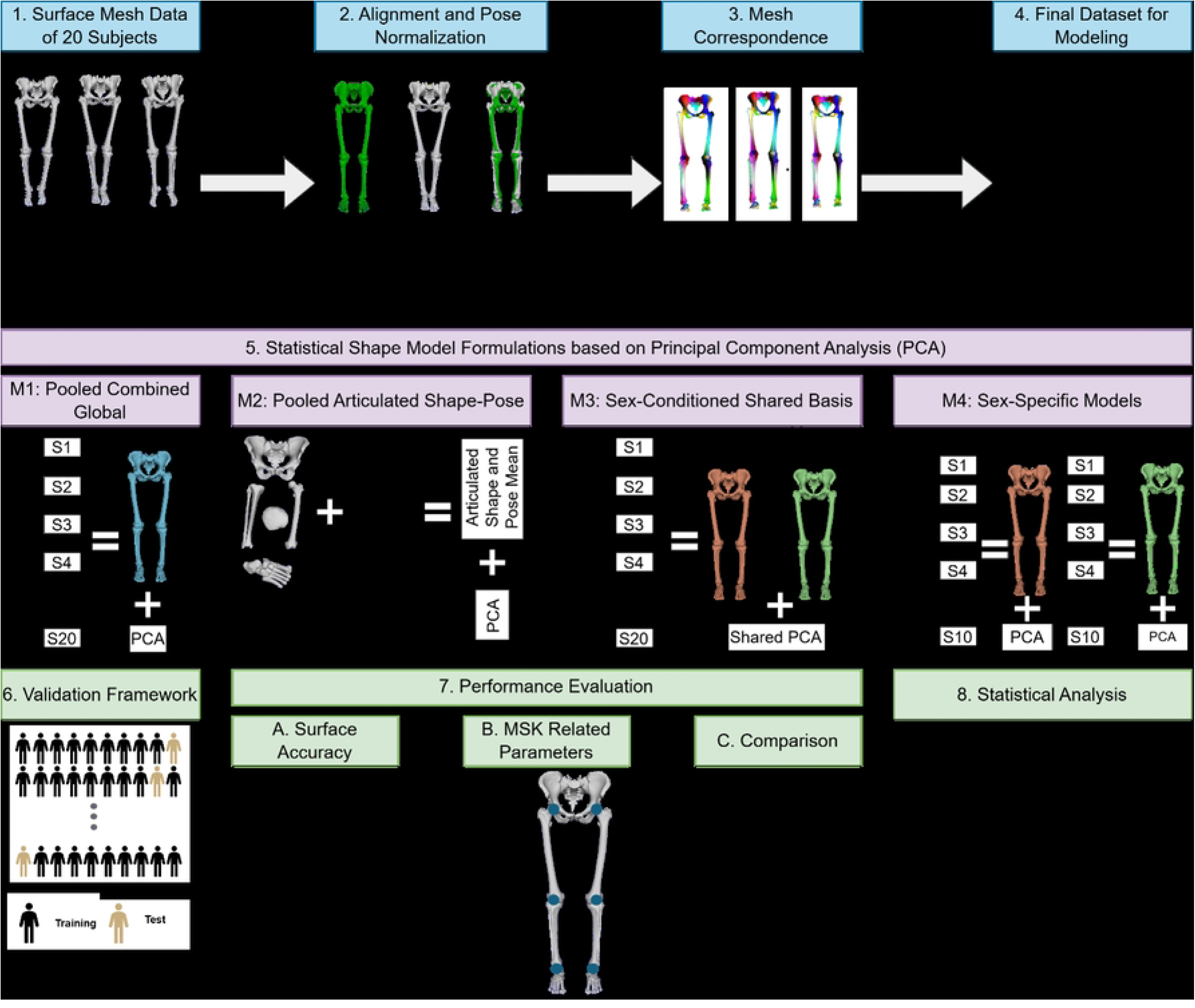
Overview of the study methodology. Surface meshes from 20 subjects were first aligned and pose-normalized to a common reference configuration, and then dense mesh correspondence was used to generate the final modeling dataset. Four principal component analysis (PCA)-based statistical shape model (SSM) formulations were then developed: M1, a pooled combined-global model; M2, a pooled articulated shape-pose model; M3, a sex-conditioned shared-basis model; and M4, separate sex-specific models. Model performance was evaluated using LOSO dense-surface reconstruction, including surface reconstruction accuracy, MSK-STAPLE-derived anatomical parameters, comparison with a study-specific anisotropic-scaling baseline, and statistical analysis.

### 2.1. Data Source

Lower-body bone geometries were obtained from the open-source dataset published by Fischer [32]. The present analysis included 20 adults with complete required anthropometric and bone data, comprising 10 female and 10 male subjects. The mean ± standard deviation, with minimum-maximum values in parentheses, for age, height, mass, and body mass index were 51.75 ± 21.48 years (19-95 years), 1.72 ± 0.08 m (1.52-1.87 m), 69.84 ± 12.53 kg (50.45-90 kg), and 23.42 ± 3.27 kg/m^2^ (17.87-28.73 kg/m^2^), respectively.

Fischer [32] obtained computed tomography (CT) scans of the subjects from the Virtual Skeleton Database (VSD) [33]. These CT scans were originally provided by the forensic institutes of the universities of Bern and Zürich, following ethical approval from the Cantonal Ethics Committee Bern [34]. The CT scans were then segmented using a semi-automatic thresholding-based algorithm. The lower threshold was set to 200 Hounsfield units (HU), while the upper threshold was set to the maximum HU value present in the volume data. This was followed by manual post-processing and, where necessary, manual segmentation to obtain bone surface files. Further information about the dataset is provided in the cited literature [32].

It should be noted that Fischer [32] reported data from 30 subjects; however, 10 subjects lacked complete anthropometric or bone data and were therefore excluded from the present study. The identification numbers of the 20 complete subjects are explicitly provided by Fischer [32]. The anatomical domains used to generate the SSMs included the pelvis, represented by the left hemipelvis, right hemipelvis, and sacrum with coccyx, as well as the bilateral femora, patellae, tibiae, fibulae, tali, and calcanei. In this study, the calcaneus domain, denoted as calcn, refers to the Rajagopal/OpenSim foot segment and includes the calcaneus together with the non-talar tarsal bones, namely the cuboid, navicular, and lateral, intermediate, and medial cuneiforms, as well as the metatarsal bones. Toe and phalangeal segments were not included in the present models. The present study used previously published data and involved no new participant recruitment or image acquisition. Ethical approval for the source CT data is reported in the original dataset publications [32–34].

### 2.2. Spatial Alignment and Pose Neutralization

The bone meshes were originally expressed in the world coordinate system (CS) of each CT scan and therefore differed in global position and articulated pose across subjects. Spatial alignment and pose normalization were performed to reduce non-anatomical positional and pose variation before correspondence registration and SSM construction.

This was achieved by first estimating the anatomical and joint CS of each bone using Shared Tools for Automatic Personalized Lower Extremity modeling (STAPLE) [35]. Briefly, the bone meshes were first pre-aligned using their principal axes of inertia. Bone-specific algorithms were then applied to identify anatomical landmarks, articular surfaces, and geometric features, such as fitted cylinders, spheres, and planes, from which CS origins and orthonormal axes were derived. The pelvic CS was based on the ASIS landmarks and ISB recommendations. The femoral CS was defined using the femoral head center and distal femoral knee geometry, while the tibial CS was aligned with the femoral knee axis and the talocrural joint center. The talar CS was derived from geometric primitives fitted to the talar trochlea and subtalar articular surfaces. The calcaneus/foot CS was defined relative to the subtalar joint frame and an auxiliary foot-sole frame, both based on calcaneus, talus, and metatarsal landmarks.

The pose of the Rajagopal OpenSim model [36] was selected as the neutral reference pose because it is widely used in MSK modeling and enables the generated SSMs to be later used to personalize OpenSim models [37]. The alignment procedure consisted of two main steps. First, a pelvis-based global rigid alignment was applied to map each subject into the Rajagopal ground CS while preserving the subject’s original articulated pose. Second, hierarchical articulated pose neutralization was performed separately for the right and left lower limbs along the kinematic chain pelvis-femur-tibia-talus-calcn/foot. At each level, the current bone frame was aligned to the corresponding Rajagopal reference frame about a joint-centered pivot, and the same correction was propagated to all downstream bones to preserve skeletal connectivity.

The patella was not used to drive the alignment; instead, it followed the corresponding femoral correction. Distal foot alignment was corrected using a pivot-preserving approach when required to avoid an unstable foot orientation. The output of this procedure was a set of pose-neutralized meshes and updated bone frames, all expressed in the Rajagopal-aligned global coordinate system. These aligned meshes were then used for MeshMonk registration. Because pose normalization used subject-specific MSK-STAPLE coordinate systems and joint-centered transformations, subsequent joint-center outcomes were interpreted relative to the same STAPLE-derived anatomical framework rather than as independent anatomical validation. The mathematical formulation of the pose-normalization procedure is provided in Section 1 of S1 Appendix.

### 2.3. Mesh Correspondence

After each bone mesh was transformed into its STAPLE-derived local CS, mesh correspondence was established by first selecting a representative template mesh for each bone. Template selection was based on shape centrality: each candidate mesh was rigidly aligned, without scaling, to the remaining subject-specific meshes; symmetric nearest-neighbor distances were then computed; and the candidate with the lowest distance-based objective score was selected as the reference template. The selected template was subsequently registered to each target mesh using MeshMonk [38]. For the pelvis, template selection and registration were performed on a single merged mesh comprising the left and right hemipelves and the sacrum. In contrast, the tibia and fibula were treated as separate bones and registered independently.

MeshMonk [38] performed an initial rigid alignment of the template to the target, followed by regularized non-rigid deformation that warped the template surface onto the target while preserving the template topology and vertex numbering. This process generated anatomically corresponding meshes with the same number and order of vertices across all subjects. For the pelvis, the merged mesh was used to preserve the spatial relationships among the left and right hemipelvis and the sacrum; however, non-rigid correspondence was constrained separately within each anatomical domain. All other bones were registered individually using conservative non-rigid settings, with surface normals and orientation-based inlier weighting enabled. The MeshMonk [38] registration settings for each anatomical domain are provided in Table B in S1 Appendix. The mean ± SD whole-body nearest neighbor distance was 0.66 ± 0.08 mm for all subjects, 0.65 ± 0.11 mm for male subjects, and 0.67 ± 0.03 mm for female subjects. Similarly, the mean bone-wise registration error was less than 1 mm for all bones.

### 2.4. Statistical Shape Modeling

After establishing mesh correspondence, principal component analysis (PCA) was used to generate SSMs. The SSMs were designed to test how different modeling choices affect dense-surface reconstruction and their potential future use in MSK-model personalization. Four model families were evaluated: a pooled combined-global model (M1), an articulated local-shape and pose model (M2), a sex-conditioned shared-basis model (M3), and separate sex-stratified models (M4).

All models used the same anatomical domains: hip_r_, hip_l_, sacrum, femur_r_, femur_l_, patella_r_, patella_l_, tibia_r_, tibia_l_, fibula_r_, fibula_l_, talus_r_, talus_l_, calcn_r_, and calcn_l_. Here, calcn_r_, and calcn_l_ refer to the Rajagopal-model calcaneus/foot segments while subscripts *l* and *r* indicate left and right sides respectively. The talus was modelled separately, and toe segments were not included in the SSMs.

#### 2.4.1. General PCA Formulation

For each model, the subject-specific representation was assembled into a subject-by-variable matrix. Let x_i_ be the vector representation of subject i. A reconstructed shape instance using k principal components was represented as:

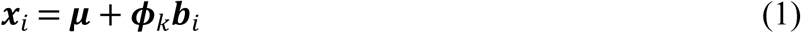

where ***μ*** is the model mean, ***φ****_k_* contains the first k principal component loading vectors, and ***b****_i_* is the vector of principal component scores. Scores were obtained by projecting the centered subject vector onto the retained PCA basis:

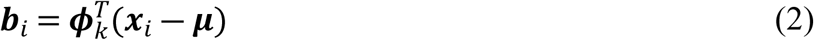

Principal component modes were visualized by deforming the mean shape by three standard deviations along each mode:

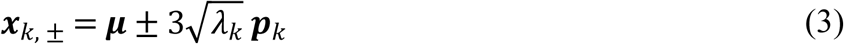

where *λ_k_* is the eigenvalue and ***p****_k_* is the loading vector of the k^th^ principal component. PCA was implemented using dual-space formulation, which is computationally efficient when the number of coordinate variables is much larger than the number of subjects [39, 40].

#### 2.4.2. Model 1: Pooled Combined Global SSM

M1 was defined as a pooled combined-global SSM. All registered bones were expressed in the common Rajagopal-aligned global CS and concatenated into one whole-lower-body vector for each subject:

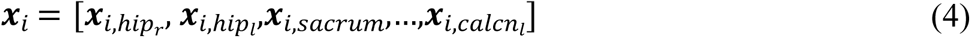

Each block contained the stacked x, y, and z coordinates of the corresponding registered mesh. This representation preserves the global spatial relationships among bones, including bilateral proportions, inter-bone positioning, residual pose variation, and covariance across the included lower-body domains.

Bone-balanced weighting was applied before PCA in M1. This correction was used to prevent larger or more densely sampled domains from dominating the covariance structure solely because they contained more vertices. For domain d with n_d_ vertices, all coordinates in that domain block were divided by the square root of n_d_ before PCA:

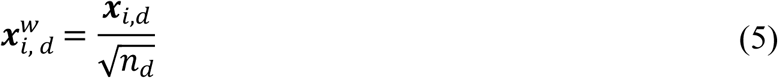

The weighted subject vectors were assembled into the matrix X^w^, and PCA was fitted in the weighted space. The exported mean shapes, reconstructed meshes, and mode shapes were converted back to physical global millimeter coordinates after PCA by reversing the block weighting. Thus, the PCA basis was learned from the bone-balanced representation, while all reported geometry and errors remained in the original physical coordinate space.

#### 2.4.3. Model 2: Articulated Local Shape and Pose SSM

M2 was defined as a pooled articulated shape-pose SSM. Unlike M1, which represented all bones directly in global coordinates, M2 separated each anatomical domain into a local bone-shape block and a local-to-global pose block. The local shape block consisted of the MeshMonk-registered mesh coordinates expressed in the STAPLE-derived local CS. The pose block described how the local mesh was positioned in the common Rajagopal-aligned global CS.

For domain d, the local-to-global transformation was:

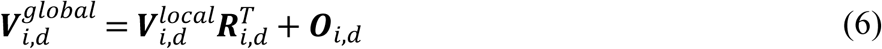

where 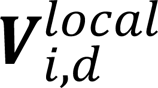 contains local registered mesh coordinates, ***R****_i_*_,*d*_ is the local-to-global rotation matrix, and ***O****_i_*_,*d*_ is the global origin of the local CS. Each domain contributed a block containing the local shape, the global origin, and the flattened rotation matrix:

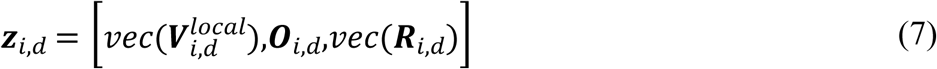

The full articulated vector was formed by concatenating ***z****_i_*_,*d*_ across all domains. During reconstruction, predicted local meshes were decoded, predicted rotation matrices were re-orthonormalized, and local meshes

were transformed back into the global CS using Equation (6). No additional block-wise standardization was applied across the local-shape, origin, and rotation blocks before PCA. Because these blocks contained quantities with different units and numerical ranges, M2 was treated as an exploratory implementation of an articulated shape-pose representation.

#### 2.4.4. Model 3: Sex-Conditioned Shared-Basis SSM

M3 used the same combined-global surface representation as M1 but introduced sex conditioning via sex-specific mean shapes and a single shared pooled PCA basis. Because this PCA basis was identical to that used in M1, M1 and M3 necessarily had identical eigenvalue and compactness spectra. The two formulations differed in the mean vector used during reconstruction. In this formulation, male and female subjects were allowed to have different baseline morphologies, while the principal modes of variation were estimated from the full cohort. This avoided the instability of fitting separate PCA bases to smaller sex-specific subsets.

Bone-balanced weighting was applied before PCA using the same domain-wise square-root weighting described in Equation (5). PCA was fitted to the pooled weighted matrix. Male and female mean vectors were then computed separately in the same weighted coordinate space, and each subject was reconstructed relative to the mean corresponding to their sex:

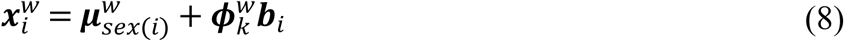

Within each LOSO fold, the male and female mean vectors were calculated using only the corresponding training subjects. After reconstruction, the weighting was removed and the result was reshaped into the full set of global bone meshes. Because M3 used the same combined-global representation as M1, no articulated decoding step was required.

#### 2.4.5. Model 4: Sex-Stratified Models

M4 used the same combined-global representation and bone-balanced weighting as M1 but fitted two independent PCA models: one using only male subjects and one using only female subjects. Each sex-specific model therefore had its own mean shape, PCA basis, scores, eigenvalues, and explained variance. This formulation tested whether fully sex-specific PCA models provided additional benefit over the sex-conditioned shared-basis formulation in M3. Because the male-only (M4-M) and female-only (M4-F) models were trained on smaller subsets, M4 was treated as an exploratory sex-stratified model.

#### 2.4.6. Virtual Joint-Center Landmarks Augmentation

In the final MSK-relevant implementation, hip, knee, and ankle joint centers derived from the neutral-pose MSK-STAPLE alignment were attached to the SSMs as virtual anatomical landmarks. These landmarks were stored as an additional three-dimensional set of points in the same Rajagopal-aligned global CS as the registered bone meshes. The virtual landmark vector for each subject was:

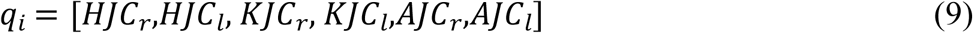

For the combined-global models that did not explicitly contain local pose origins (M1, M3, and M4), q_i_ was appended to the weighted surface vector as an additional anatomical block:

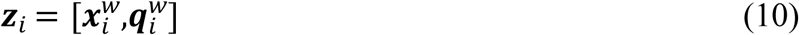

where 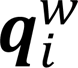 was obtained by dividing the virtual-landmark block by the square root of six, corresponding to the six virtual joint-center points. This gave the joint center block a controlled contribution to the PCA while preserving the original surface representation. A unit block gain was used in the final analysis. After PCA reconstruction, the virtual landmark block was transformed back to physical millimeter coordinates and used to compute joint-center errors and hip joint-center width.

During LOSO evaluation of M1, M3, and M4, the held-out subject’s STAPLE-derived reference joint centers were excluded from coefficient estimation. Each fold was trained using the surfaces and virtual landmarks of the training subjects. Coefficients for the held-out subject were estimated from its registered surface block, and the corresponding joint-center block was reconstructed from the retained PCA representation. Thus, this analysis evaluated cross-block reconstruction of STAPLE-derived joint centers from a dense surface representation.

M2 was handled differently because its articulated pose block already contained the same STAPLE-derived origins. The femur origins corresponded to the HJCs, the tibia origins to the KJCs, and the talus origins to the AJCs. These origins were verified to match the exported MSK-STAPLE joint centers to numerical precision. Therefore, separate virtual joint-center landmarks were not appended to M2; joint centers were instead extracted directly from the reconstructed pose-origin terms. Because these pose origins were variables within the articulated representation, the M2 joint-center errors quantify reconstruction of represented pose-origin terms rather than fully independent joint-center prediction.

### 2.5. Model Comparison

The SSMs were first summarized by compactness, defined as the number of principal components required to explain 99% of the training-set variance within each model’s representation. Because the formulations used different feature spaces, compactness values were interpreted within rather than directly across model types. Generalization was evaluated using LOSO reconstruction. For each fold, the model was trained on the remaining subjects, and coefficients for the held-out subject were estimated from its complete registered surface representation. The retained PC count was selected independently within each training fold as the number required to explain 99% of that fold’s variance. Accordingly, this LOSO analysis quantified dense-surface representational generalization and not reconstruction from sparse markers or anthropometric measurements.

#### 2.5.1. Difference Metrics

Reconstruction performance of SSMs was evaluated against the original geometries using whole-body, bone-wise, and MSK-related error metrics. The definitions of these metrics are provided as follows:

Surface generalization accuracy was compared using original-mesh nearest-neighbor RMSE. Original meshes were used as reference surfaces because they lacked dense vertex correspondence with the reconstructed SSM meshes. For the reconstructed vertex *j* on the bone *b*, the nearest-neighbor distance was defined as:

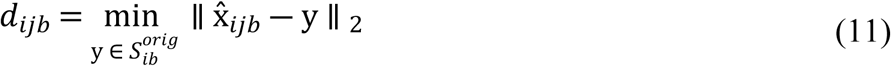

where x̂*_ijb_*is the reconstructed SSM vertex, and 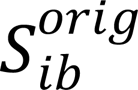 is the original surface of bone *b* for the subject *i*. Bone-wise nearest-neighbor RMSE was calculated as:

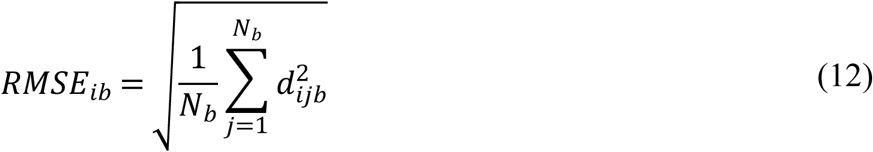

where *N_b_*is the number of reconstructed vertices for the bone *b*. Whole-body RMSE was calculated by pooling the squared nearest-neighbor distances across all included bones:

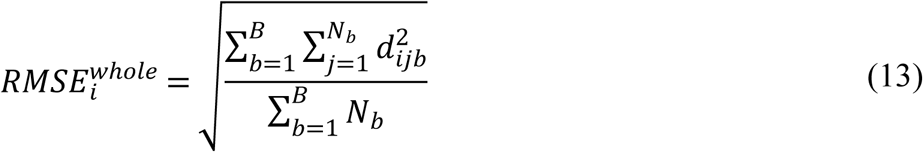

where *B* denotes the set of included bones. The primary surface outcomes were whole-body LOSO RMSE and bone-wise LOSO RMSE, both expressed in millimeters. Distances were calculated in one direction, from each reconstructed vertex to its nearest neighbor on the corresponding original mesh. Consequently, this metric may depend on mesh sampling and was interpreted as a reconstruction-error measure rather than a symmetric surface-distance metric.

MSK-relevant reconstruction accuracy was evaluated relative to the bilateral joint-center coordinates obtained from MSK-STAPLE. For the joint center *J*, where *J* ∈ [*HJC*,*KJC*,*AJC*], and side *s* ∈ [*L*,*R*], the side-specific Euclidean error was:

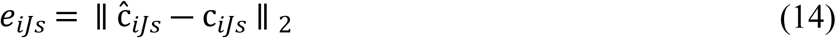

where c*_iJs_*is the reference joint center, and ĉ*_iJs_*is the model-reconstructed joint center. The bilateral mean joint-center error was calculated as:

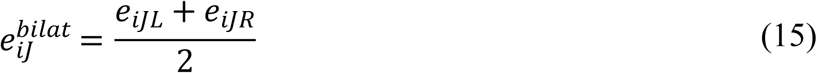

The bilateral mean error was used for reporting and statistical analysis. Hip joint-center width was defined as the distance between the left and right hip joint centers:

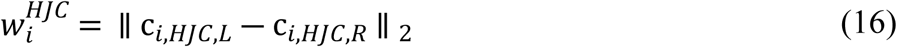

and the HJC-width error was:

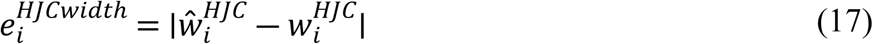

Thus, the retained MSK outcomes were bilateral HJC, KJC, and AJC errors, as well as HJC-width error.

#### 2.5.2. Comparison of SSMs with Each Other

For pairwise comparisons between SSMs, paired error differences were computed for each subject as:

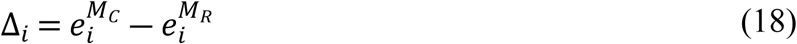

where 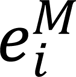 denotes the error for the subject *i* using the model *M*, *M_C_* is the comparison model, and *M_R_* is the reference model. The error metric *e* is either whole-body RMSE, bone-wise RMSE, bilateral joint-center error, or HJC-width error, depending on the comparison. Negative values indicated lower error for the comparison model than for the reference model.

#### 2.5.3. Comparison of SSMs with Anisotropic Linear Scaling

In addition to the comparisons among SSMs, each SSM was compared with a study-specific pelvis-referenced bounding-box anisotropic-scaling baseline. For each held-out subject, the training-set mean shape was scaled to the subject’s shape using the subject’s bounding-box dimensions. Independent scale factors were computed along the global coordinate axes as:

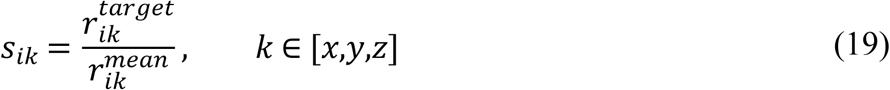

where 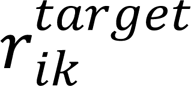 is the held-out subject’s lower-body bounding-box range along the axis *k*, and 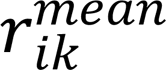 is the corresponding range of the LOSO training-set mean. Using these scaling factors, the same transformation was applied to both surface vertices and joint-center coordinates about the pelvis body-CS center.

For SSM-versus-scaling comparisons, paired differences were calculated as:

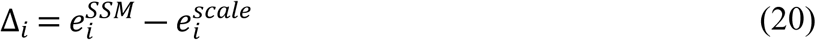

where *e* is either whole-body RMSE, bone-wise RMSE, bilateral joint-center error, or HJC-width error. Negative values indicated that the SSM had lower error than anisotropic linear scaling. This comparator provided a geometric reference for the present dataset and was not intended to reproduce a complete marker-based OpenSim scaling workflow. Because the SSM coefficients and scaling factors were estimated using different amounts of subject-specific geometric information, the comparison should not be interpreted as a matched-input evaluation of clinical personalization methods.

### 2.6. Statistical Analysis

All primary statistical analyses were performed using subject-level LOSO reconstruction errors. Model compactness was reported descriptively as the number of PCs required to explain 99% of the training-set variance and was not subjected to inferential testing. Statistical testing was restricted to planned contrasts defined a priori from the model-comparison framework.

For comparisons among SSMs, paired differences were computed using the SSM-versus-SSM paired-difference metric defined above. In the all-subject analysis, M2 and M3 were compared with the pooled reference model M1. Because M4 was fitted as separate sex-specific models, M4 was evaluated only within the corresponding sex-stratified subsets. Within females, M4-F was compared with M1, M2, and M3. Within males, M4-M was compared with M1, M2, and M3. For all SSM-versus-SSM comparisons, negative paired differences indicated lower error for the comparison model than for the reference model.

For comparisons with the pelvis-referenced anisotropic linear-scaling baseline, paired differences were computed using the SSM-versus-scaling paired-difference metric defined above. Each SSM was compared with the scaled mean shape within the relevant analysis group: M1, M2, and M3 for all subjects; M1, M2, M3, and M4-F for females; and M1, M2, M3, and M4-M for males. Negative differences indicated lower error for the SSM than for linear scaling.

Because each subject contributed matched errors across models or reconstruction methods, all planned comparisons were tested using two-sided paired Wilcoxon signed-rank tests. Tests were performed for whole-body RMSE, bone-wise RMSE, bilateral HJC, KJC, AJC, and HJC-width errors. LOSO reconstruction was used as the primary analysis to evaluate subjects excluded from model training, while each held-out subject’s registered surface remained available for coefficient estimation.

For whole-body RMSE and MSK-related parameter errors, p-values from the planned paired Wilcoxon tests were adjusted using the Holm correction within each analysis group and outcome family. For bone-wise RMSE, the same planned paired Wilcoxon tests were performed separately for each bone, and p-values were adjusted using the Benjamini-Hochberg false-discovery-rate correction across the planned bone-wise comparisons. The same correction strategy was applied to both SSM-versus-SSM and SSM-versus-scaling analyses. Statistical significance was defined as an adjusted p-value less than 0.05. Given the small cohort and the strongly overlapping LOSO training sets, inferential results were interpreted as exploratory and considered together with the magnitude and direction of the paired differences.

ChatGPT 5.6 Sol (OpenAI) was used during development of the computational workflow to assist with improving selected MATLAB and Python code. All AI-assisted code changes were reviewed by the authors, revised where necessary, and subsequently executed and tested before being used in the reported analyses. The authors take full responsibility for the final code, analyses, and results.

## 3. Results

### 3.1. First Three Principal Components

The mode-shape visualizations in Fig. 2 showed that PC1 produced the largest displacement from the mean shape, consistent with PC1 explaining the greatest proportion of variance. Based on directional inspection of the mode shapes, PC1 primarily captured superior-inferior variation, reflecting longitudinal or height-related changes in the lower-limb skeleton. PC2 was dominated by anterior-posterior variation, while PC3 mainly represented mediolateral variation. These directions should be interpreted as dominant trends rather than pure single-axis motions, because each PC also included coupled regional shape changes across the skeleton. PC2 and PC3 showed smaller and more localized effects, particularly around the pelvis, knee, ankle, and foot regions. The distal lower-limb structures were repeatedly affected across modes. M1 and M3 displayed similar deformation patterns, whereas M2 showed more pronounced distal and articulated variation, consistent with its shape-pose formulation. The sign of each PC is arbitrary; therefore, the +3 SD and -3 SD shapes should be interpreted as opposite directions along the same mode.

**Fig. 2.**
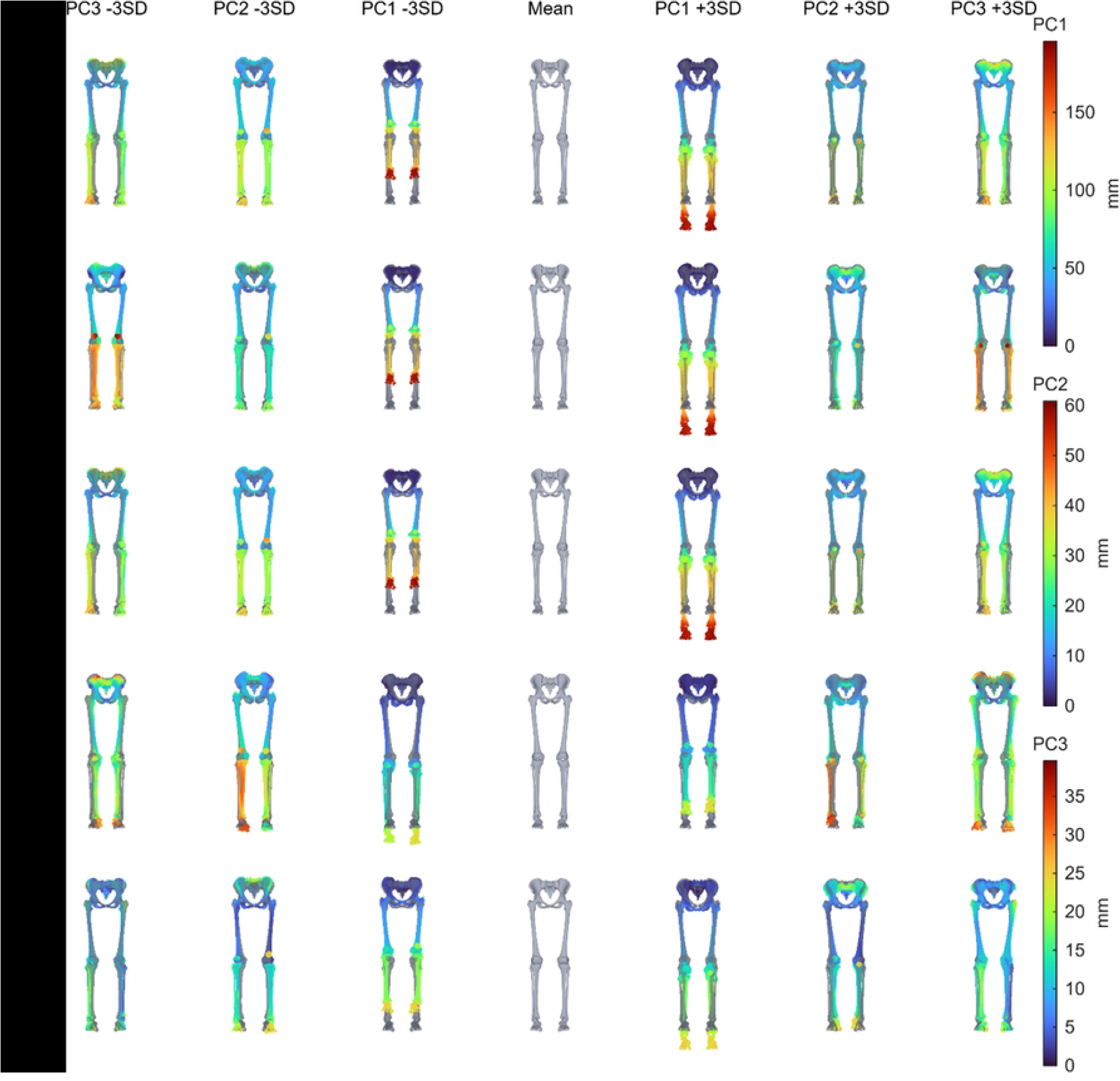
Mode-shape visualization of the first three principal components (PCs) for each statistical shape model (SSM). Rows correspond to the different SSMs, and columns show the mean shape and the shapes reconstructed at ±3 standard deviations (SD) along PC1, PC2, and PC3. Surface heat maps show the magnitude of displacement from the corresponding model mean, in millimeters, highlighting the anatomical regions most affected by each mode. The mean shape is shown in grey. Separate color scales are used for PC1, PC2, and PC3; therefore, displacement magnitudes should be compared within each PC rather than directly across PCs.

### 3.2. Model Compactness

Model compactness, expressed as the variance explained by individual PCs, cumulative variance, and the number of PCs required to explain 99% of the total variance, is shown in Fig. 3. For all SSMs, PC1 accounted for the largest proportion of variance, ranging from 62.47% in M2 to 90.16% in M1 and M3, while each subsequent PC accounted for less than 15% of the variance. M1 and M3 shared the same pooled PCA basis and therefore had identical compactness spectra by construction: PC1 explained 90.16% of the total variance, and nine PCs were required to explain 99% of the variance in both models. M2 distributed variance across a different feature space: PC1 explained 62.47% of the variance, and 17 PCs were required to explain 99%. This difference may reflect its articulated representation containing both local bone-shape and pose parameters; consequently, compactness was interpreted within rather than directly across the different model formulations. The sex-specific SSMs also captured more variance in PC1 than M2, with PC1 explaining 87.87% and 73.64% of the variance in M4-F and M4-M, respectively. Within the sex-specific PCA spaces, 7 PCs were required to explain 99% of the variance for M4-F, while 8 PCs were required for M4-M. These values were interpreted within each sex-specific model because M4-F and M4-M were fitted to smaller subgroup datasets and therefore had fewer available PCs than the full-cohort models. During LOSO validation, the reconstruction threshold was selected separately within each training fold as the number of PCs required to explain 99% of that fold’s training-set variance.

**Fig. 3.**
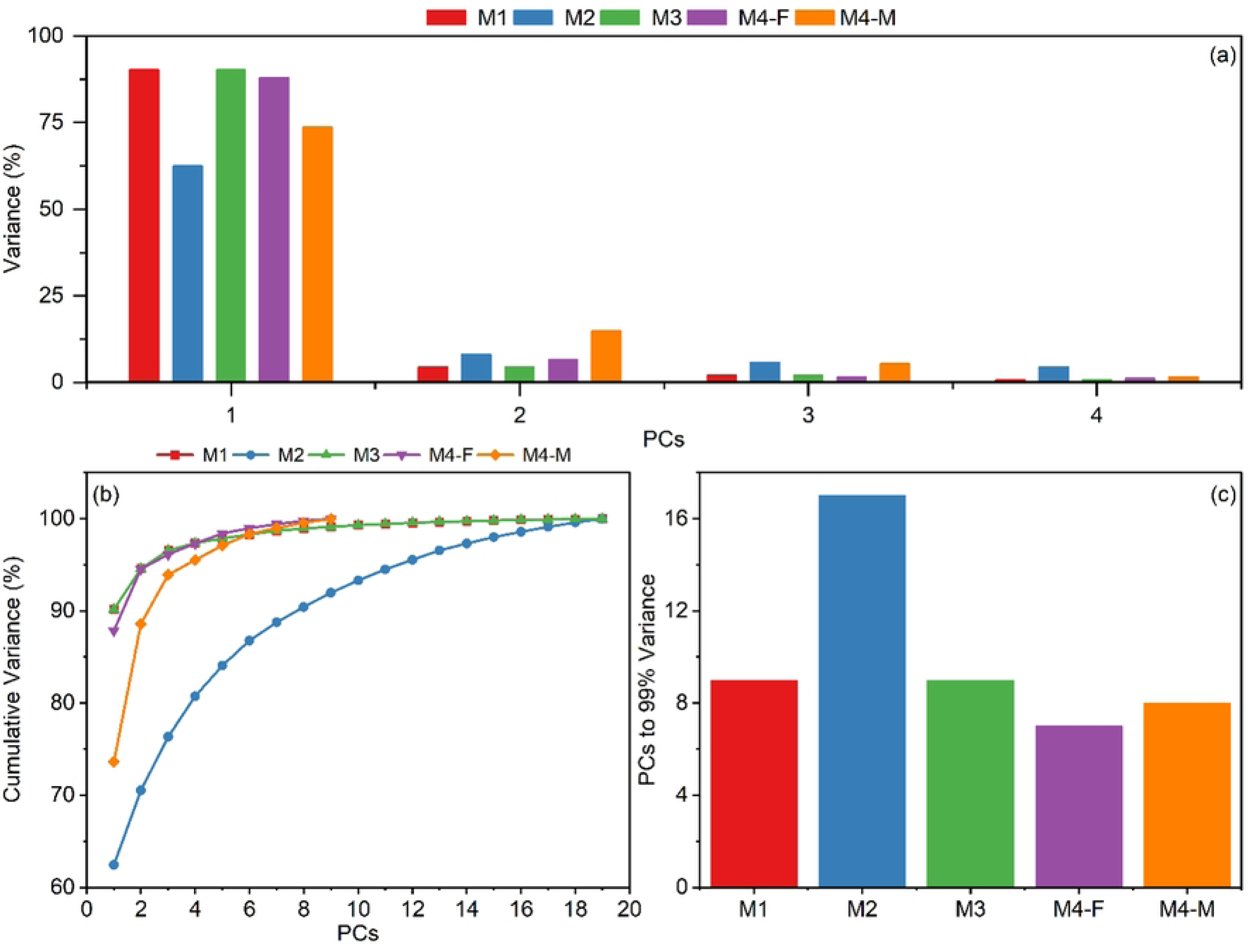
Variance explained by the principal components (PCs) of each statistical shape model (SSM). (a) Percentage of variance explained by each individual PC. (b) Cumulative variance explained as a function of the number of PCs for each SSM. (c) Number of PCs required to explain 99% of the total variance for each SSM.

### 3.3. Whole Body LOSO Generalization

Fig. 4 summarizes the LOSO reconstruction accuracy of the SSMs, including whole-body surface RMSE, planned model comparisons, joint-center errors, and HJC-width errors. Among the full-cohort models, M1 and M3 had the lowest whole-body reconstruction errors, whereas M2 produced consistently higher RMSE. M2 exceeded M1 by 1.63 mm, and this difference was significant after Holm adjustment (p < 0.001), while M3 differed from M1 by only -0.01 mm and was not significantly different (p = 0.411). Within the present cohort, the independently fitted M4 models did not show lower whole-body reconstruction error than M1 or M3. M4-F had significantly higher RMSE than M1 and M3 (Holm-adjusted p = 0.029 for both) but did not differ significantly from M2. M4-M had significantly higher RMSE than M1 and M3 (Holm-adjusted p = 0.012 for both) but had lower RMSE than M2 by 0.93 mm (Holm-adjusted p = 0.037).

**Fig. 4.**
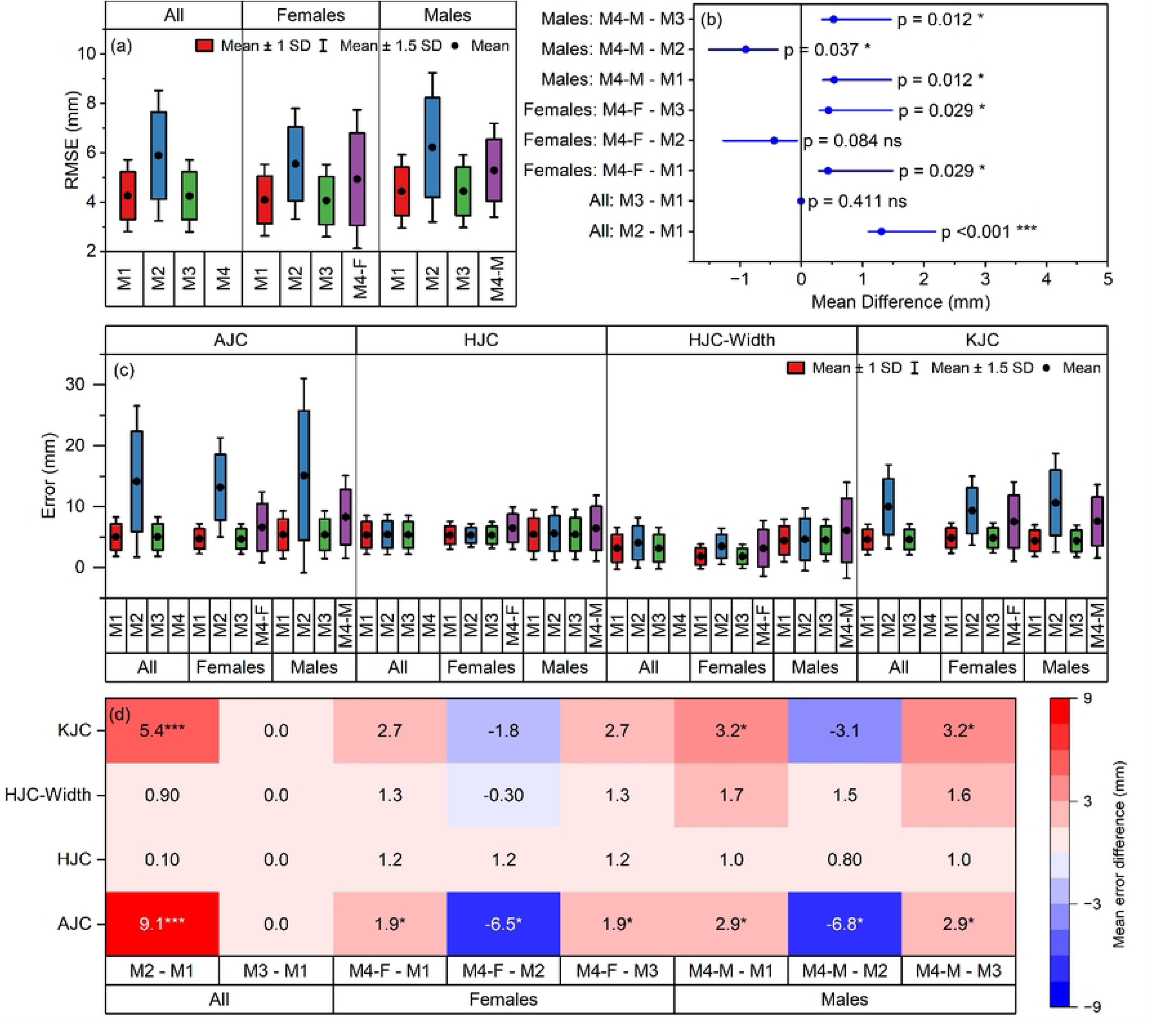
Leave-one-subject-out (LOSO) reconstruction accuracy of the statistical shape models (SSMs). (a) Whole-body surface reconstruction RMSE for all subjects, female subjects, and male subjects. (b) Planned pairwise comparisons of whole-body RMSE among the SSMs, reported as mean differences calculated as comparison model minus reference model, with Holm-adjusted p-values. (c) LOSO errors in anatomical parameters, including bilateral ankle joint center (AJC), hip joint center (HJC), knee joint center (KJC), and hip joint center width (HJC-width), computed between reconstructed SSM outputs and the corresponding target anatomy. (d) Heat map of planned pairwise mean differences in joint-center and HJC-width errors. Statistical significance was based on Holm-adjusted p-values and is indicated as follows: ns, adjusted p ≥ 0.05; *, adjusted p < 0.05; **, adjusted p < 0.01; and ***, adjusted p < 0.001. Here, ns denotes non-significant.

Joint-center analysis showed that the largest model-dependent differences occurred at the ankle and knee. Compared with M1, M2 increased AJC and KJC errors by 9.1 mm and 5.4 mm, respectively, and both differences were significant. In contrast, M3 showed small mean differences from M1 across the evaluated anatomical parameters. The between-model mean differences were smaller for HJC and HJC-width errors, generally remaining ≤1.7 mm. For females, M4-F had significantly higher AJC error than M1 by 1.9 mm (Holm-adjusted p = 0.039), whereas its HJC, HJC-width, and KJC differences from M1 were not significant (1.2 mm, p = 0.252; 1.3 mm, p = 1.000; and 2.7 mm, p = 0.059, respectively). For males, M4-M had significantly higher AJC and KJC errors than M1 by 2.9 mm (p = 0.041) and 3.2 mm (p = 0.029), respectively, whereas its HJC and HJC-width differences were not significant (1.0 mm, p = 0.393; and 1.7 mm, p = 1.000, respectively). Overall, no statistically significant HJC or HJC-width differences were detected in the planned comparisons; this should not be interpreted as evidence of equivalence. AJC error was significantly higher for both sex-specific models, while KJC error was significantly higher for M4-M.

### 3.4. Bone-wise LOSO Generalization

To further interpret the LOSO findings, bone-wise reconstruction errors are presented in Fig. 5. This analysis shows that the higher whole-body RMSE observed for M2 was mainly driven by distal bones rather than by uniform increases across the skeleton. Compared with M1, M2 produced significantly higher errors in the tibiae, tali, fibulae, and calcanei, with mean RMSE differences of 2.57-3.33 mm for the tibiae, 4.28-4.48 mm for the tali, 2.79-3.02 mm for the fibulae, and 3.20-4.11 mm for the calcanei. In contrast, M2 showed lower errors for the sacrum and hip bones, indicating that its reduced performance was primarily associated with the distal limb and hindfoot domains. M1 and M3 showed very similar bone-wise performance, and no statistically significant differences were detected for the individual bones, consistent with their similar whole-body LOSO accuracy. The sex-specific M4 models showed slightly higher bone-wise errors than M1 and M3 in several bones, although these RMSE differences were generally not significant at the individual-bone level. This suggests that the higher whole-body RMSE observed in the sex-specific models was distributed across multiple domains rather than driven by a single anatomical region. Overall, the bone-wise analysis indicates that M1 and M3 provided the most consistent reconstruction accuracy, while the main limitations of M2 were associated with greater variability in distal lower-limb bones.

**Fig. 5.**
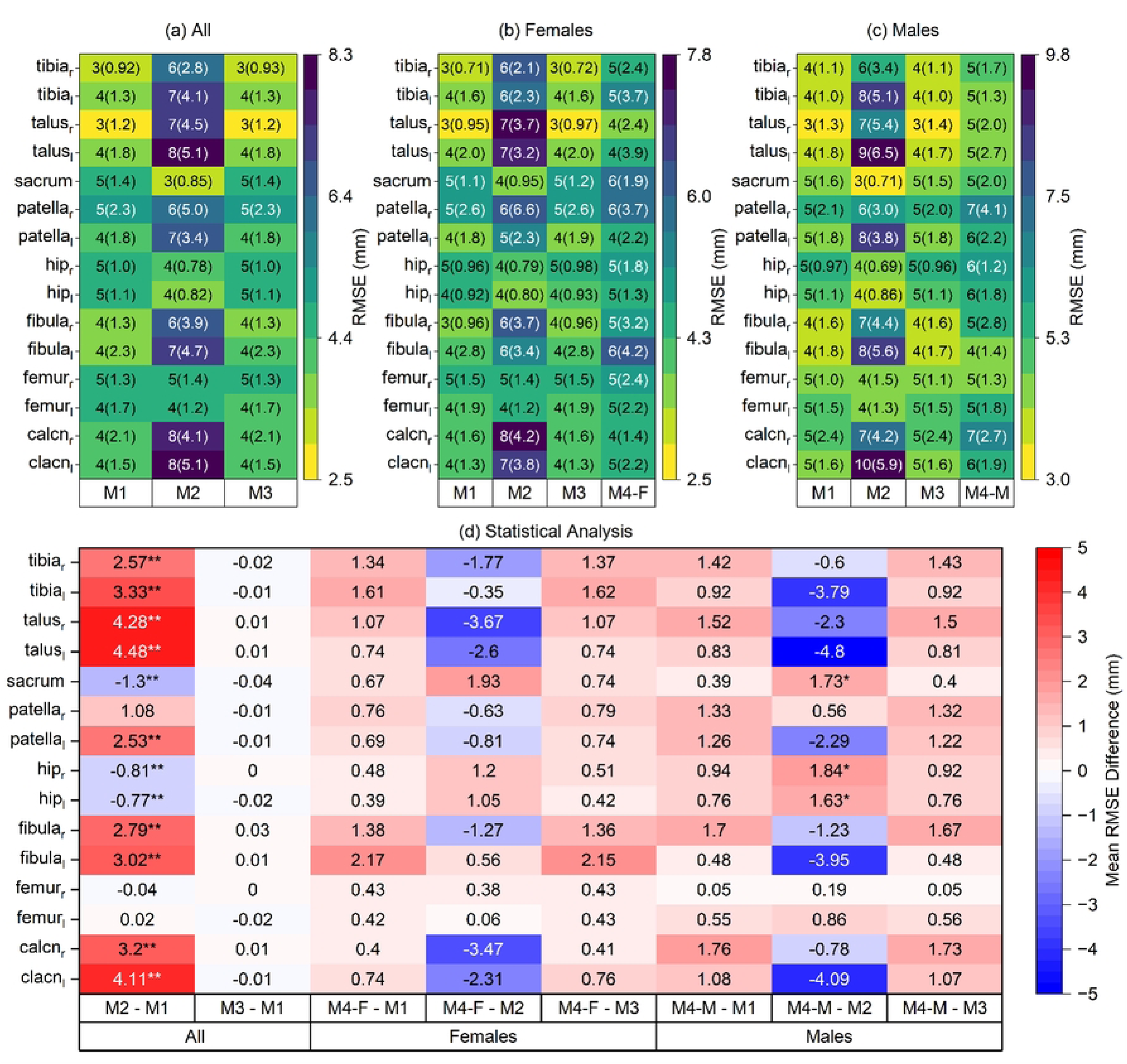
Bone-wise leave-one-subject-out (LOSO) reconstruction performance of the statistical shape models (SSMs) for (a) all subjects, (b) female subjects, and (c) male subjects. Heat-map cells report mean RMSE, with SD in parentheses, in millimeters for each bone and model. Panel (d) shows the planned pairwise comparisons of bone-wise RMSE, reported as mean differences calculated as comparison model minus reference model. Positive values indicate higher error for the comparison model, whereas negative values indicate lower error. Statistical significance was based on Benjamini-Hochberg false-discovery-rate-adjusted p-values and is indicated as follows: ns, adjusted p ≥ 0.05; *, adjusted p < 0.05; **, adjusted p < 0.01; and ***, adjusted p < 0.001.

The spatial error maps in Fig. 6 support the bone-wise LOSO findings. M1 and M3 showed broadly similar error distributions, with relatively low-to-moderate errors across most of the skeleton. In contrast, M2 showed visibly larger errors in the distal lower limb, particularly around the ankle, talus, calcaneus, and distal tibia/fibula regions, consistent with the higher bone-wise RMSE observed for the tibiae, tali, fibulae, and calcanei. Across models, the most challenging regions to reconstruct were the ankle–hindfoot complex, distal tibia/fibula, and localized pelvic and knee regions. The long-bone shafts generally showed lower errors, suggesting that reconstruction difficulty was concentrated near complex joint surfaces, small bones, and regions with higher anatomical curvature. The sex-specific models showed more spatially variable error patterns, with localized increases in selected pelvic and distal-limb areas. Overall, these maps indicate that higher RMSEs of M2 were primarily driven by distal lower-limb reconstruction errors, whereas M1 and M3 provided more spatially consistent reconstructions across the articulated skeleton.

**Fig. 6.**
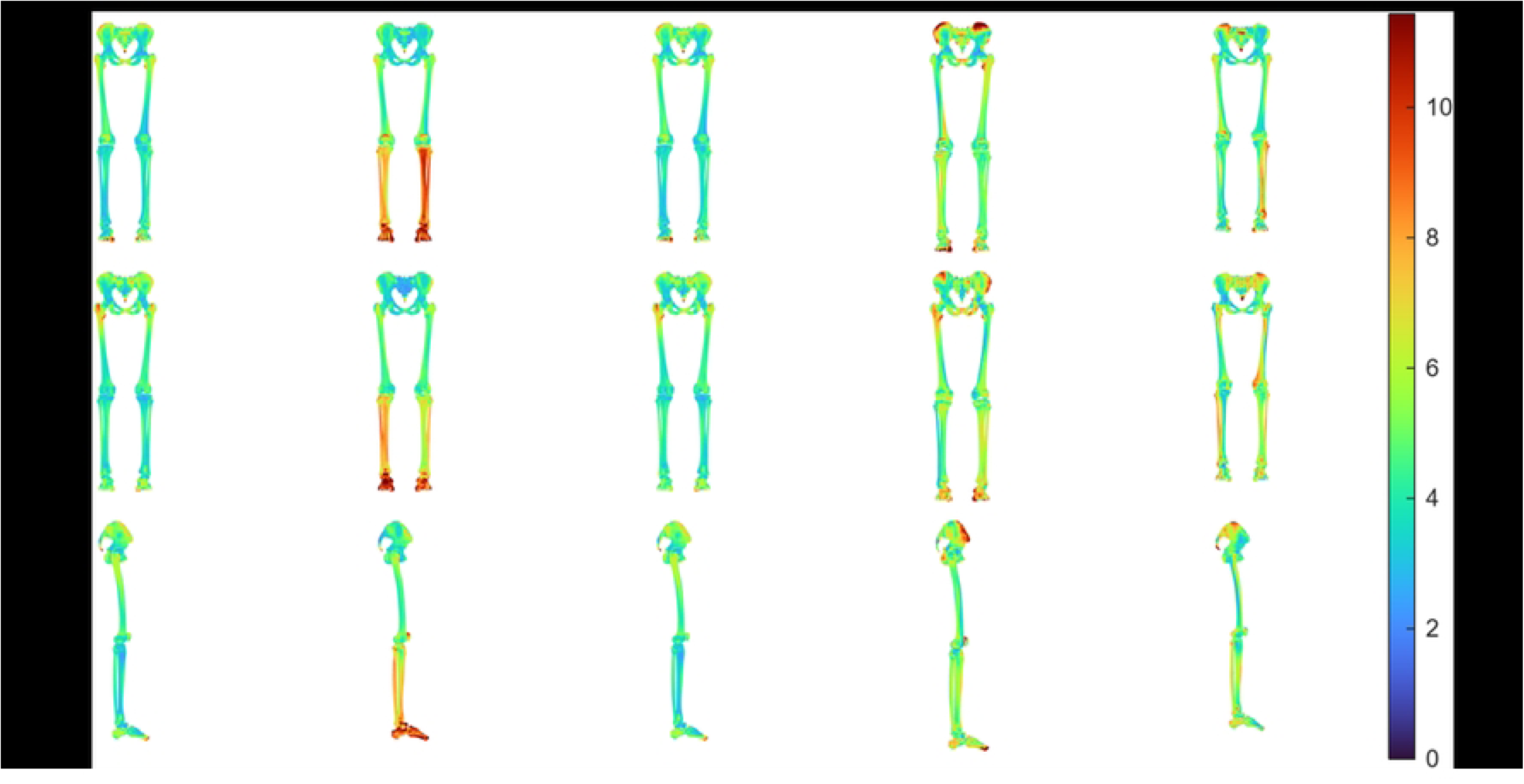
Spatial distribution of leave-one-subject-out (LOSO) reconstruction RMSEs for the statistical shape models (SSMs). Columns show the reconstructed error maps for M1, M2, M3, M4-M, and M4-F, respectively. Rows show anterior, posterior, and lateral views of the lower-limb skeleton. Surface colors represent the magnitude of reconstruction error in millimeters, with warmer colors indicating larger errors.

### 3.5. Comparison with Linear Scaling

The comparison between SSM-based LOSO reconstruction and the anisotropic linear-scaling baseline is summarized in Fig. 7. Within the present dense-surface LOSO framework, all SSMs had significantly lower whole-body surface RMSE than the study-specific anisotropic-scaling baseline. For all subjects, the mean RMSE reductions were approximately 4.1 mm for M1, 2.5 mm for M2, and 4.0 mm for M3 (Holm-adjusted p < 0.001). Similar improvements were observed in the sex-specific analyses, with whole-body RMSE reductions ranging from approximately 2.5 to 4.3 mm across the female and male comparisons. Bone-wise comparisons showed that the largest improvements generally occurred in the distal lower limb, particularly the tibiae, tali, fibulae, and calcanei, where several models reduced RMSE by approximately 5-7 mm relative to anisotropic scaling. Anatomical-parameter errors were also lower with SSMs, especially at the ankle and knee joint centers. For all subjects, M1 reduced AJC and KJC errors by 12.5 mm and 11.0 mm, respectively, while M3 showed comparable reductions of 12.3 mm and 10.8 mm. Thus, under the present dense-surface reconstruction conditions, the SSMs had lower surface and STAPLE-referenced joint-center errors than the selected geometric scaling baseline, with the largest differences occurring in the distal lower limb and at the ankle and knee joint centers.

**Fig. 7.**
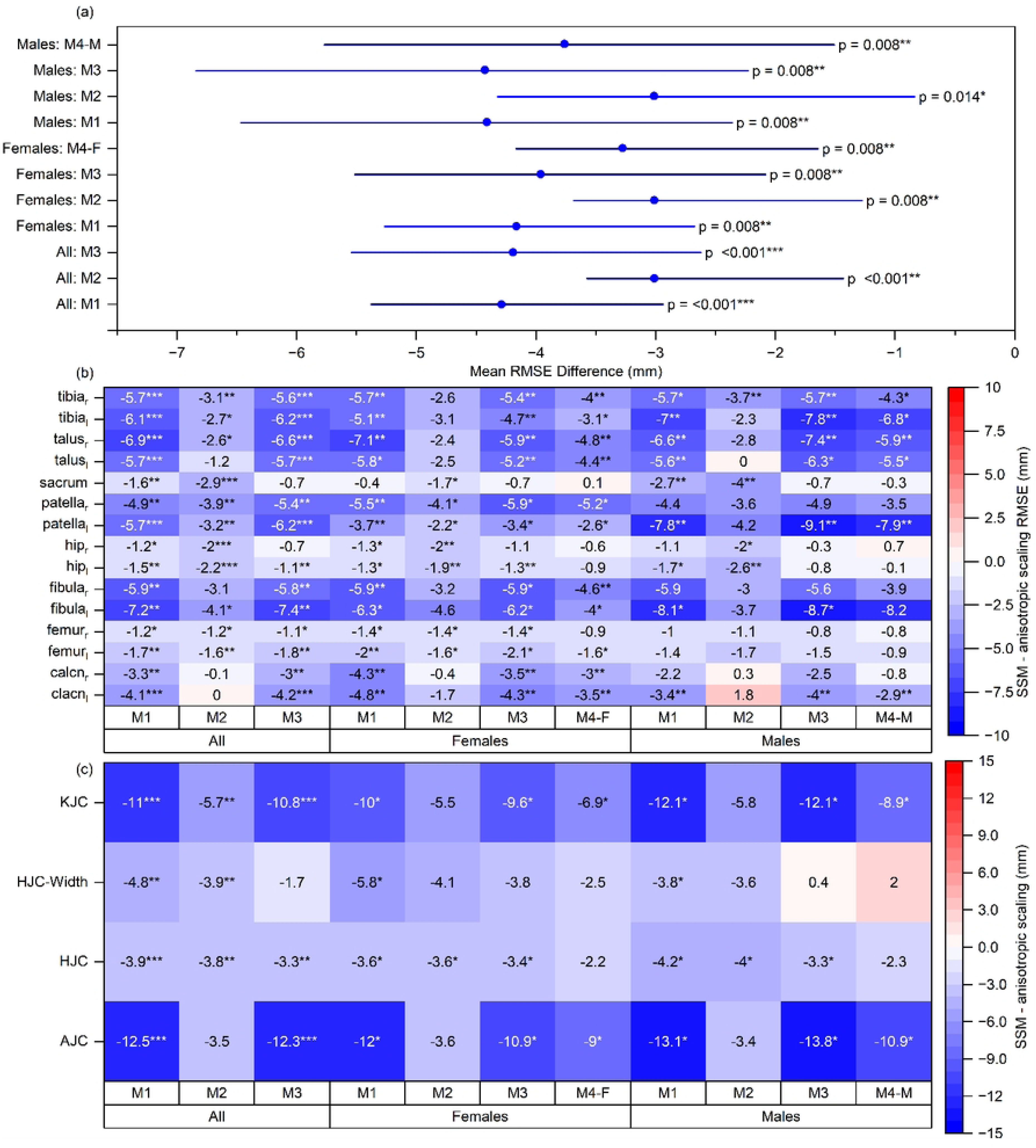
Comparison of leave-one-subject-out (LOSO) reconstruction accuracy between each statistical shape model (SSM) and the study-specific pelvis-referenced bounding-box anisotropic-scaling baseline. (a) Whole-body surface RMSE differences for all subjects, female subjects, and male subjects. (b) Bone-wise surface RMSE differences between each SSM and anisotropic scaling. (c) Differences in anatomical-parameter errors, including mean bilateral ankle joint center (AJC), hip joint center (HJC), knee joint center (KJC), and hip joint center width (HJC-width). Differences were calculated as SSM error minus anisotropic-scaling error. Negative values indicate lower error for the SSM, whereas positive values indicate lower error for anisotropic scaling. Holm-adjusted p-values were used for whole-body and MSK outcomes, and Benjamini–Hochberg false-discovery-rate-adjusted p-values were used for bone-wise outcomes. Statistical significance is indicated as follows: ns, adjusted p ≥ 0.05; *, adjusted p < 0.05; **, adjusted p < 0.01; and ***, adjusted p < 0.001. The SSM and scaling methods used different amounts of subject-specific geometric information; consequently, this represents comparison with the selected geometric baseline rather than a matched-input evaluation of clinical scaling.

## 4. Discussion

This study developed and compared four lower-body SSM formulations using dense-surface LOSO reconstruction. Within this framework, M1 and M3 produced the most consistent generalization, whereas M2 produced larger errors, particularly for distal lower-limb and STAPLE-referenced joint-center outcomes. All SSM formulations also had lower errors than the study-specific bounding-box anisotropic-scaling baseline. These findings characterize the implemented representations and comparator within the present dataset; they do not constitute a matched-input validation of sparse-marker or anthropometric personalization.

The dominance of the first principal component (PC1) was consistent with previous lower-limb SSM studies. In the present models, PC1 accounted for the largest proportion of variance, ranging from 62.47% in M2 to 90.16% in M1 and M3, while subsequent PCs each accounted for less than 15%. This agrees with Carman et al. [29], who reported that PC1 captured most retained-size variance in pediatric pelvis, femur, and tibia/fibula models, and with Zhang et al. [24], who reported that PC1 mainly represented bone size. M2 required 17 PCs to explain 99% of the variance within its articulated feature space. Because this representation combined local-shape, origin, and rotation variables with different units and numerical ranges, its variance spectrum should be interpreted within that parameterization rather than as a direct measure of anatomical complexity relative to the other models.

M1 and M3 showed the most consistent reconstruction performance. Because they shared the same pooled PCA basis, their identical compactness spectra were expected by construction. M3 differed by using sex-specific mean shapes during reconstruction. It did not show lower whole-body error than M1 in the present analysis, but it maintained similar accuracy while allowing sex-specific baseline morphologies. Consequently, M3 should be viewed as an alternative conditional formulation rather than as demonstrably preferable to M1.

Within the implemented parameterization, M2 produced the highest errors among the pooled models. Because M2 combined local-shape, origin, and rotation variables, its performance may reflect the selected feature representation, the relative numerical scaling of the feature blocks, sample size, and retained-PC threshold. The present findings should therefore not be interpreted as demonstrating a general limitation of articulated SSMs.

The sex-specific M4 models should also be interpreted cautiously. M4-F and M4-M reduced errors relative to anisotropic scaling, but neither outperformed M1 or M3 in whole-body LOSO reconstruction. M4-M did outperform M2 at the whole-body level, but it remained less accurate than M1 and M3. The smaller sex-specific training subsets likely reduced the stability of the independently fitted PCA bases. HJC and HJC-width errors were not significantly different from those of M1, whereas AJC error was significantly higher for both M4-F and M4-M, and KJC error was significantly higher for M4-M. These findings show that independently fitted sex-specific bases did not improve reconstruction in the present sample. Because each M4 LOSO fold contained only nine sex-matched training subjects, the study was not designed to determine whether sex-specific modeling would become advantageous in a larger cohort.

The bone-wise and spatial error analyses showed that reconstruction difficulty was not uniform across the skeleton. The largest model-dependent errors occurred in the distal lower limb, particularly the tibiae, fibulae, tali, calcanei, and ankle-hindfoot region. M2 increased LOSO RMSE relative to M1 by approximately 2.6-4.5 mm in the tibiae and tali and by approximately 3.2-4.1 mm in the calcanei, explaining its poorer whole-body performance and larger AJC errors. Similar challenges have been reported previously: Zhang et al. [24] observed local errors up to 15 mm in some regions, and Nolte et al. [22] noted peak reconstruction errors around the malleoli. These findings suggest that small, highly curved, joint-critical structures are more difficult to reconstruct than long-bone shafts.

The improvement over linear scaling is consistent with previous literature. However, direct quantitative comparison is limited because these previous studies commonly fitted SSMs from sparse landmarks, whereas the present SSM coefficients were estimated from complete registered surfaces. Zhang et al. [24] reported that an articulated adult lower-limb SSM reduced surface RMSE from 5.22 mm with isotropic scaling to 4.28 mm, while Nolte et al. [20] showed that SSM reconstructions from skin- or bone-based landmarks were more accurate than linear scaling. Carman et al. [28] similarly reported that pediatric articulated SSMs outperformed adult and pediatric linear scaling. In the present study, all SSMs reduced whole-body LOSO RMSE relative to anisotropic scaling, with mean reductions of approximately 4.1 mm for M1, 2.5 mm for M2, and 4.0 mm for M3.

The practical contribution of this study is a reproducible comparison of statistical representations across a bilateral lower-body dataset that includes distal hindfoot and metatarsal-containing foot segments. Evaluating joint-center reconstruction alongside surface errors helps characterize anatomical features relevant to future musculoskeletal model personalization. The differences observed at the ankle and knee motivate further matched-input studies to determine whether improvements in anatomical reconstruction translate into changes in kinematics, muscle moment arms, or joint loading [23].

To support reuse, LowerBodySSM v1.0.0 packages the MATLAB and Python workflow, source and derived geometries, registered meshes, statistical model datasets, validation outputs, environment information, and documentation. The interactive Companion supports workflow navigation, execution instructions, status checks, and inspection of selected outputs; long-running analyses are executed separately in MATLAB and Python. The archive has been deposited as a Zenodo draft record with reserved DOI 10.5281/zenodo.22253723. It is accessible through a private review link during peer review and will be made publicly accessible upon publication. Component-specific licenses and third-party requirements are documented within the archive.

This study has several limitations. The cohort was small, particularly for M4, and LOSO evaluation was confined to the same source dataset. The held-out subject’s complete registered surface was used to estimate SSM coefficients, whereas the scaling baseline used only three bounding-box dimensions; the comparison was therefore not input matched and should not be interpreted as a direct evaluation of clinical scaling workflows. The joint-center references and pose-normalization frames were both derived using MSK-STAPLE, so the joint-center outcomes represent internal reconstruction relative to this framework rather than independent anatomical validation. Surface accuracy was quantified using a one-directional nearest-neighbor RMSE, and the fixed 99% threshold retained nearly all available modes for some formulations. Sex-related findings may also reflect subgroup sample size and differences in body size or other demographic variables. External validation, matched sparse-input fitting, alternative surface metrics and model-complexity sensitivity analyses, and evaluation of downstream MSK outputs remain necessary.

Overall, M1 and M3 provided the most consistent dense-surface LOSO reconstruction within the current dataset. The results support their use as reference formulations for the released resource, while matched-input and external validation will be required before concluding that they improve practical MSK-model personalization over established scaling workflows.

## 5. Conclusions

This study developed a reproducible workflow for pose-normalized adult lower-body statistical shape models extending from the pelvis to the metatarsal-containing foot segments. The pooled global and sex-conditioned shared-basis formulations provided the most consistent dense-surface reconstruction within the present cohort. The sex-conditioned model did not demonstrate a whole-body reconstruction advantage over the pooled global model, while distal ankle and hindfoot structures remained challenging. These findings characterize the evaluated representations and geometric scaling comparator; they do not establish improved clinical personalization. LowerBodySSM and its interactive Companion provide a documented resource for further evaluation using larger cohorts, sparse anatomical inputs, independent joint-center references, and downstream musculoskeletal simulations.

## Data and Code Availability

The data and author-generated code supporting this study are deposited in the LowerBodySSM v1.0.0 archive in Zenodo under the reserved DOI 10.5281/zenodo.22253723. The archive includes source and derived geometries, registered meshes, statistical shape model datasets, subject-level validation outputs, result tables and figures, MATLAB and Python code, the interactive Companion, environment information, and documentation. The record is currently unpublished and will be made publicly accessible upon publication of the article. Source anatomical geometries were obtained from the dataset described by Fischer [32]. Third-party materials and their derivatives remain subject to their applicable licensing terms. Component-specific licenses, attribution requirements, and reuse conditions are documented within the archive.

## Author Contributions

Conceptualization: Muhammad Abdullah

Data Curation: Muhammad Abdullah and Abdul Aziz Hulleck

Funding Acquisition: Marwan El-Rich

Investigation: Muhammad Abdullah and Abdul Aziz Hulleck

Methodology: Muhammad Abdullah, Abdul Aziz Hulleck, and Marwan El-Rich

Supervision: Marwan El-Rich and Kinda Khalaf

Validation: Muhammad Abdullah and Abdul Aziz Hulleck

Visualization: Muhammad Abdullah

Writing – original draft: Muhammad Abdullah

Writing – review and editing: Muhammad Abdullah, Abdul Aziz Hulleck, Kinda Khalaf, and Marwan El-Rich

## Supporting Information

S1 Appendix. Supplementary methods and registration settings. Detailed pose-normalization equations, symbol definitions, and MeshMonk registration settings for the anatomical domains included in the statistical shape models.

